# Vestibular modulation of multisensory integration during actual and vicarious tactile stimulation

**DOI:** 10.1101/576611

**Authors:** Sonia Ponzo, Louise P. Kirsch, Aikaterini Fotopoulou, Paul M. Jenkinson

## Abstract

**Background:** The vestibular system has been shown to contribute to multisensory integration by balancing conflictual sensory information. It remains unclear whether such modulation of exteroceptive (e.g. vision), proprioceptive and interoceptive (e.g. affective touch) sensory sources is influenced by epistemically different aspects of tactile stimulation (i.e. felt from within vs seen, vicarious touch).

**Objective:** We aimed to i) replicate previous findings regarding the effects of galvanic stimulation of the right vestibular network (i.e. LGVS) in multisensory integration and ii) examine vestibular contributions to multisensory integration when touch is felt but not seen (and vice-versa).

**Method:** During artificial vestibular stimulation (LGVS, RGVS and Sham), healthy participants (N=36, Experiment 1; N=37, Experiment 2) looked at a rubber hand while either their own unseen hand or the rubber hand were touched by affective or neutral touch.

**Results:** We found that i) LGVS led to enhancement of vision over proprioception during visual only conditions (replicating our previous findings), and ii) LGVS (vs Sham) favoured proprioception over vision when touch was felt (Experiment 1), with the opposite results when touch was vicariously perceived via vision (Experiment 2), and with no difference between affective and neutral touch.

**Conclusions:** We showed how vestibular signals modulate the weight of each sensory modality according to the context in which they are perceived and that such modulation extends to different aspects of tactile stimulation: felt and seen touch are differentially balanced in multisensory integration according to their epistemic relevance.

**Highlights:**

- LGVS increased proprioceptive drift during vision of a rubber hand
- Touch on participant’s hand decreased proprioceptive drift during LGVS
- Vicarious touch on the Rubber Hand increased proprioceptive drift during LGVS
- Vestibular signals differently balance sensory sources in multisensory integration

## Introduction

During multisensory integration, signals from different sensory modalities are weighted according to their contextual reliability and combined to produce a unitary perceptual experience of the world [1–3] and our own body, including the sense of body ownership (i.e. the sense that our body belongs to us; [4]). Body ownership has been extensively studied using the Rubber Hand Illusion (i.e. RHI; [5]), during which participants watch a fake hand being touched in- or out-of-synchrony with their own unseen hand. The RHI has provided significant insight into how conflict between vision (of a synchronously touched rubber hand) and proprioception (of the real hand’s location) is resolved by information from one modality (vision) dominating over the others (proprioception and touch; [6–8]). This “visual capture” effect (i.e. dominance of visual cues over other modalities; [9]) occurs, in particular, when vision is deemed contextually most reliable (e.g. when we process visuo-proprioceptive signals in the horizontal plane; [6]). However, not all sensory conflictual situations are solved with a dominance of vision over proprioception. For example, an illusory feeling of movement is often experienced whilst sitting on a stationary train and observing an adjacent train beginning to move past. In such instances, the vestibular system, primarily involved in regulating balance and coordination during self-motion, also contributes to multisensory integration, providing information signalling an unresolved conflict between vision (i.e. “I see motion”) and proprioception (i.e. “I feel I am *not* moving”).

Clinical and experimental studies further suggest that the vestibular system plays an important role in the multisensory integration processes contributing to body ownership [10–19], although findings were not always consistent (e.g. see Galvanic Vestibular Stimulation, i.e. GVS [20], studies [21,22]). A previous study from our group [23] aimed to clarify these conflicting findings, whilst assessing how vestibular and interoceptive signals (i.e. feelings about the physiological condition of one’s own body; [24,25]) interact to shape body ownership. Recent research indicates that body ownership is modulated by interoceptive signals [26,27], and can be enhanced by applying gentle touch at slow velocities that activate specialised nerve fibres (CT afferents), which provide interoceptive information in the form of tactile pleasure [28–32]. Administering GVS during a RHI procedure using slow, affective (CT-afferent-optimal) or fast, emotionally neutral (CT-afferent sub-optimal) touch, we found that right-hemisphere vestibular stimulation increased proprioceptive drift during “vision-only” (i.e. pure visual capture) and synchronous visuo-tactile conditions (with no observable effect on subjective embodiment) [23]. Moreover, the enhancement of proprioceptive drift during right vestibular stimulation was greater following affective compared with neutral touch conditions. These findings were interpreted as a right-hemisphere stimulation-induced enhancement of vision over proprioception (see also [33,34]). However, the specific mechanism by which touch enhances body ownership during right-hemisphere vestibular stimulation remains unclear. Given that affective touch has been shown to elicit comparable feelings of pleasure, as well as neural activation in the posterior insula, both when experienced directly on one’s own skin and when observed on someone else’s skin (i.e. vicarious affective touch; [35,36]), the contribution of affective touch to body ownership may not only depend on its felt components but also on its *seen*, vicarious aspects. Thus, the vestibular system may differentially modulate such contribution according to the way touch is perceived.

The current work sought to address this outstanding ambiguity, by dissociating felt and seen touch during two RHI experiments with concurrent GVS. We included conditions during which slow (affective) or fast (neutral) touch was applied only to the real hand without concurrent touch on the rubber hand (Experiment 1), and vice-versa (Experiment 2), to determine whether the enhancement of proprioceptive drift was driven by the *seen* or the *felt* component of affective touch. Specifically, we aimed to i) replicate our previous findings [23], showing that vision of a rubber hand during right-hemisphere vestibular stimulation leads to increased proprioceptive drifts towards the rubber hand, even without touch (“visual capture of proprioception”) and ii) explore whether vestibular stimulation would favour proprioception over vision when touch is felt but not seen, but favour vision over proprioception when touch is seen but not felt. We hypothesised that administering affective touch only on participant’s own hand during stimulation of the right vestibular network would lead to smaller proprioceptive drifts (i.e. disruption of a previously induced “visual capture”), compared with neutral, fast touch, whilst affective touch on the rubber hand only would have opposite effects (i.e. enhancement of “visual capture”) due to the vicarious properties of affective touch. We did not expect to observe changes in embodiment questionnaire scores, since all touch conditions in the current study involved visuo-tactile asynchrony, consistently found not to elicit increased embodiment feelings [37,38].

## Materials and methods

### Participants

In Experiment 1, thirty-six, right-handed, healthy participants (23 females, age range=18-48 years, M=24.39; SD=6.01), were recruited via an institutional subject pool. Two participants were excluded from the analysis (they scored more than 2.5 SD away from the mean in more than 2 distributions). The final sample consisted of 34 participants (22 females; age range=18-48 years, M=24.24, SD=5.90). Thirty-seven new healthy participants (24 females, age range: 18-44, M=22.27; SD=5.35 years), partook in Experiment 2. Two participants were excluded (see above for criteria), with a final sample of 35 participants (22 females; age range=18-44, M=22.26; SD=5.44).

Exclusion criteria included psychiatric/neurological history, vestibular disturbances, pregnancy or metal plaques in participants’ body and previous participation in GVS studies (due to the necessary deception involved in sham conditions). Both studies were approved by an institutional Ethics Committee and all participants gave written consent.

### Experimental design

We applied GVS (LGVS, RGVS, Sham) during a RHI task in a within-subjects, block design, with the order of the three GVS blocks counterbalanced across the sample. Each of the 3 GVS blocks comprised two stroking conditions (slow, affective at 3 cm/s or fast, neutral at 18 cm/s touch [29], administered in a counterbalanced order across participants), each preceeded by a visual only condition (pure “visual capture”) (see Figure 1A and 1B). Stroking conditions (Figure 1C) were administered to explore whether touch on participant’s hand only during LGVS would reduce a previously induced visual capture, with the prediction that affective touch would have a further disrupting effect than neutral touch.

**Figure 1.**
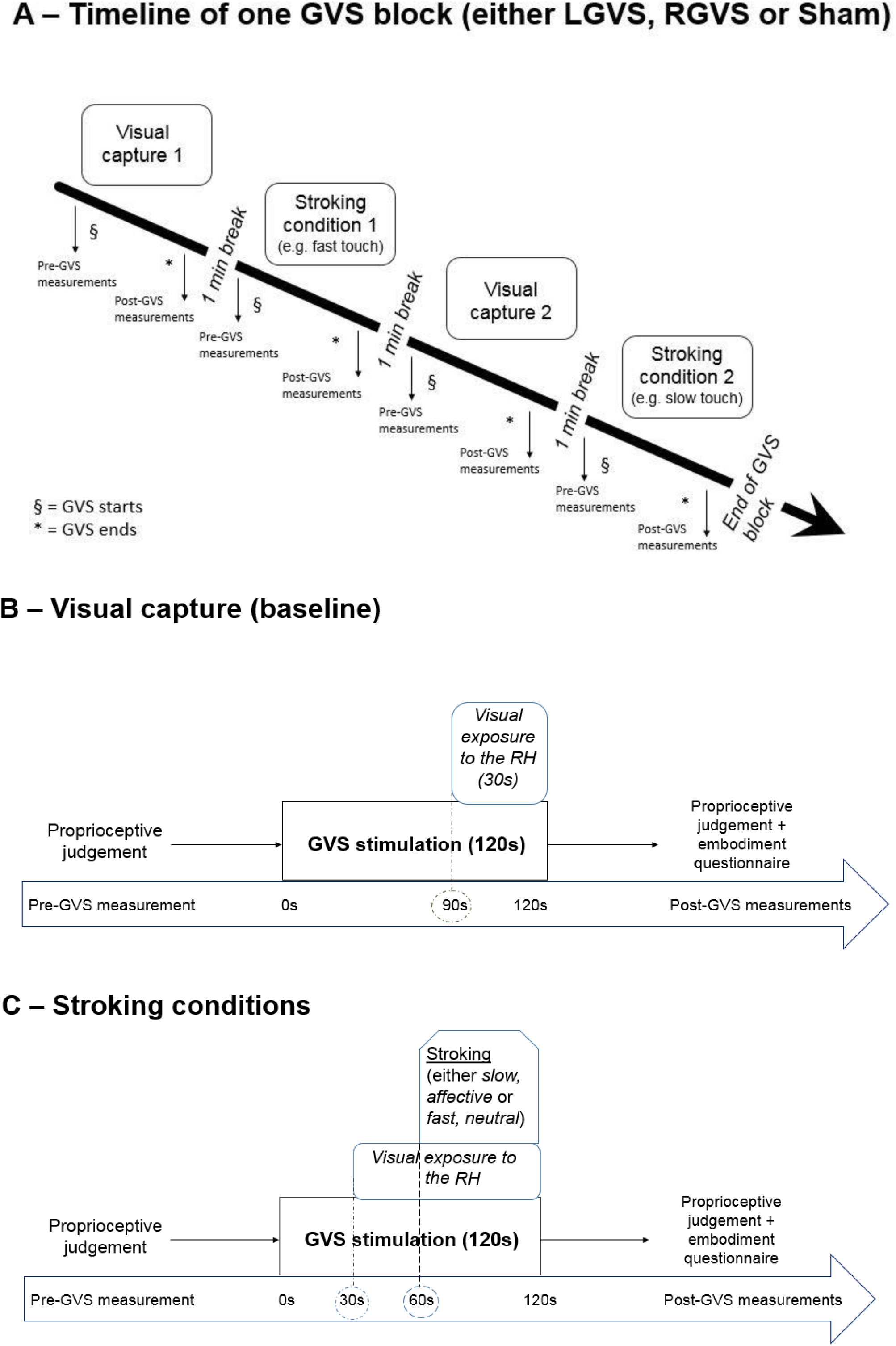
A) *Timeline of one prototypical GVS block.* At the beginning of each of the three GVS blocks (either LGVS, RGVS or Sham), participants undertook the first visual capture baseline, with measures taken before and after stimulation (see B for details). Subsequently, one of the two stroking conditions (see C for further details) was conducted in a counterbalanced order, with measures taken before and after stimulation. B) *Timeline of the visual capture baselines*. Before the visual capture condition started, participants performed a proprioceptive judgement (pre-GVS measurement). Immediately afterwards, the vestibular or sham stimulation commenced, lasting for 2 minutes, during which participants sat with their eyes open. During the last 30 seconds of vestibular or sham stimulation, the experimenter opened the box lid and instructed the participant to look at the rubber hand until told otherwise. After 120 seconds (total) stimulation the lid was closed and participants immediately performed a second proprioceptive judgement and completed the embodiment questionnaire (post-GVS measurements). C) *Timeline of the stroking conditions*. Both stroking conditions (slow, affective or fast, neutral touch) followed the same structure. Participants made an initial (pre-GVS measurement) proprioceptive judgement, followed immediately by vestibular or sham stimulation lasting for 120 seconds. After 30 seconds of vestibular stimulation, the rubber hand was revealed by the experimenter and participants were asked to continuously look at it for 30 seconds. Then, the experimenter started stroking participants (experiment 1) or rubber hand’s (experiment 2) forearm slowly or fast for 60 seconds, while the participant was asked to keep looking at the rubber hand. At the end of the 2 minutes, both tactile and vestibular stimulation ended and participants were asked to perform a second proprioceptive judgement and answer the embodiment questionnaire (post-GVS measurements).

Two outcome measures were collected: proprioceptive drift (i.e. the perceived shift of the participant’s hand towards the rubber hand, in centimetres) and an Embodiment Questionnaire [23]. Proprioceptive drift was assessed pre-GVS and post-GVS for each condition and calculated by subtracting the post-GVS estimate of the left hand’s location from the pre-GVS one (Fig. 1.A). At the end of each condition, participants completed the Embodiment Questionnaire [39], presented on a computer in a randomised order (see Section 1, Supplementary Materials). The answers to each question were averaged in order to obtain an overall embodiment score per condition.

The design and procedure for Experiment 2 were identical to Experiment 1 except for the fact that touch was applied on the Rubber Hand only (instead of participant’s own hand) and that we predicted an enhancement rather than a disruption of visual capture effects.

### Experimental setup and materials

#### Galvanic Vestibular Stimulation

We implemented a bipolar stimulation with fixed intensity (1mA) and duration (2 minutes per condition), delivered via a direct current stimulator (NeuroConn DC-stimulator, neuroCare Group GmbH, München, Germany). The total amount of stimulation per GVS block was 8 minutes, with each experiment involving 24 minutes (including Sham) of non-continuous stimulation. Each GVS block was followed by a 20-minute break in order to minimise possible stimulation after-effects [20].

GVS was delivered via two 3×3cm carbon rubber electrodes fixed either on the participants’ mastoid bones (LGVS and RGVS) or neck (Sham) using a rubber band. During LGVS (i.e. left-anodal/right-cathodal stimulation), the anode was on the left mastoid process and the cathode on the right. During RGVS (i.e. left-cathodal/right-anodal), the inverse configuration was used. During Sham, the electrodes were placed on the nape (∼5 cm below the end of the mastoid processes).

#### Rubber Hand Illusion

The apparatus was the same as detailed in our previous study [23], with the exception of the distance between the rubber and participant’s hand (see Figure 2).

**Figure 2.**
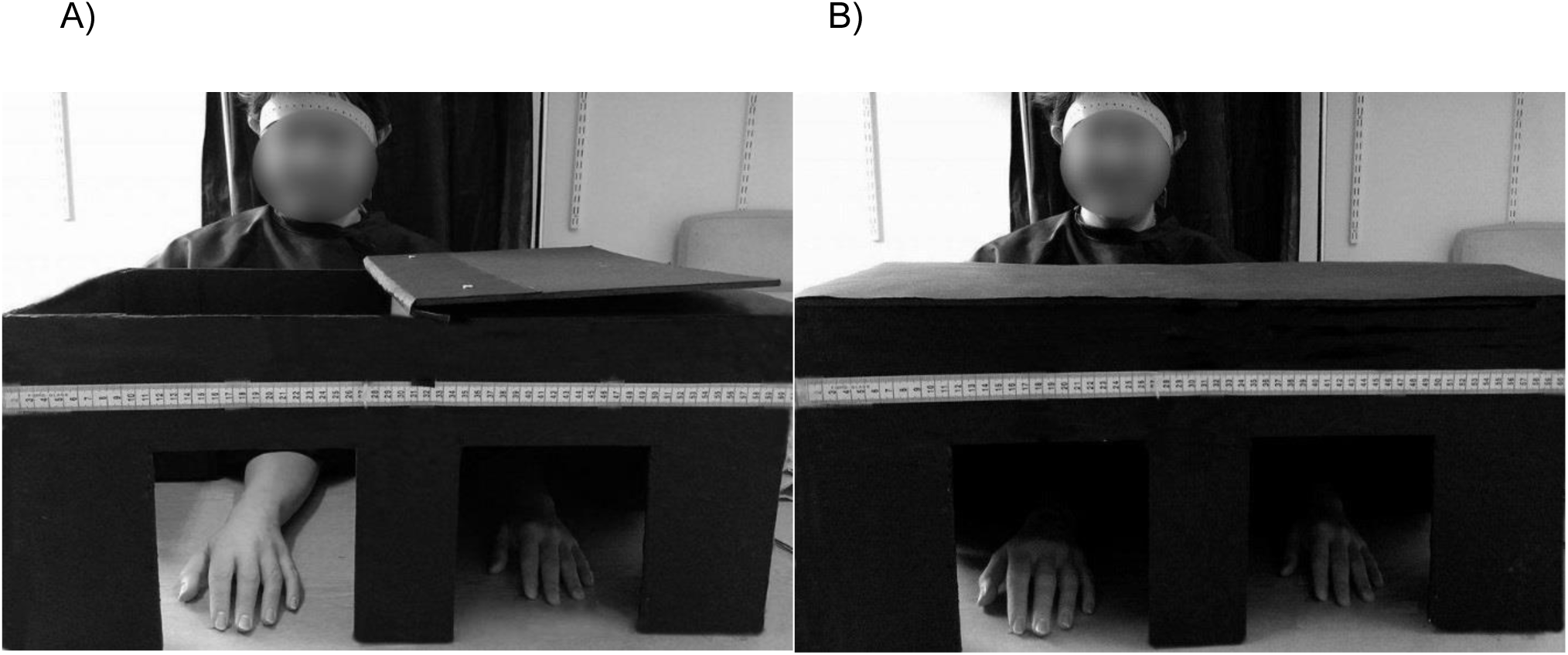
Participants were asked to place their left hand into the left half of the box, while the rubber hand was positioned in the right half, in line with the centre of participant’s torso (distance between the rubber hand and the participant’s hand=27.5 cm). Both the participant’s and rubber hand’s left index fingers were located on the marked spots. A) While the box was open, participants were asked to look inside and observe the rubber hand. B) Before and after each condition, with the box closed and covered by a black carton, participants had to perform a proprioceptive judgement (see Experimental procedure).

### Experimental Procedure

Participants positioned their left forearm in the box and the experimenter aligned their index finger with the rubber hand (hidden from participant’s view). Each condition started with a proprioceptive judgement (pre-GVS measurement; see [23]), followed by GVS stimulation. During the first condition (visual capture, Figure 1B), the rubber hand was only revealed after 1 minute and 30 seconds of stimulation and participants were asked to continuously look at the rubber hand for the last 30s. Participants then performed a second proprioceptive judgement with the box closed and completed the embodiment questionnaire (post-GVS measurements).

After the first visual capture condition, there was a one-minute break, during which participants were asked to move their left arm to reduce any cumulative effects. During the break (in Experiment 1 only), two adjacent 9×4cm areas were drawn with a washable marker on the participant’s left forearm, to control for stroking pressure and habituation [29]. Subsequently, one of the two stroking conditions began (slow or fast velocity), with a pre-GVS proprioceptive judgement (Figure 1C). Immediately afterwards, the vestibular stimulation started and for the first 30s participants sat without performing any task. Then, the experimenter opened the lid and asked participants to focus on the rubber hand. After 30s of visual only exposure to the rubber hand, the experimenter started stroking the participants’ forearm (Experiment 1) or the Rubber Hand (Experiment 2), either slowly at 3 cm/s (i.e. single touch=3s) or fast at 18 cm/s (i.e. single touch=0.5s) for 60s. Touch was always administered proximally to distally and each stroke was followed by a 1s pause. Participants were instructed to maintain their gaze on the rubber hand. After 120s, the vestibular-tactile stimulation ended, the post-GVS proprioceptive judgement was obtained and participants answered the embodiment questionnaire (post-GVS measurements). The second visual capture and stroking conditions of the block began after a one-minute break.

### Manipulation checks

At the end of the experiment, with no vestibular stimulation applied, participants were asked to rate the pleasantness of two sets of stroking (at 3cm/s and 18cm/s) to ensure that they perceived slow touch as more pleasant than fast touch [23,28–30]. In Experiment 2, we included an extra block of trials in which we asked participants to rate pleasantness of strokes observed on the rubber hand only (the vicarious and actual stroking block were counterbalanced - see SM, section 2.2.5.), to check whether they would rate the seen affective touch as more pleasant than the seen neutral touch. This was confirmed by parametric and non-parametric tests (see SM, 2.1.4 and 2.2.4). Participants were then asked to report any physical sensation associated with the vestibular stimulation and guess in which of the three configurations they thought they had received vestibular stimulation (SM, 2.1.5. and 2.2.5.).

### Data analysis

We first aimed to replicate previous findings of increased visual capture following LGVS [23] in both experiments. We conducted i) a 3 (GVS: LGVS vs. RGVS vs. Sham) × 2 (Order: first vs. second visual capture) repeated-measures ANOVA on the proprioceptive drifts obtained in the two separate visual capture conditions, to check for variability over time and ii) a one-way (GVS: LGVS vs. RGVS vs. Sham) ANOVA on the averaged proprioceptive drift values of the two visual capture conditions. Since we hypothesised a right-hemisphere-specific effect of vestibular stimulation on visual capture, we conducted planned contrasts between LGVS and Sham, and between LGVS and RGVS using Bonferroni-corrected paired-samples t-tests (α=0.025). Additionally, we checked whether combining the proprioceptive drift values obtained in both experiments (N=69) following the two visual capture conditions in the three GVS blocks (as well as their averages) would lead to the same effects found in the two separate experiments. In order to do so, we applied the same aforementioned analyses.

In Experiment 1, we aimed to examine the effects of vestibular stimulation and touch on visual capture. We performed separate 3 (GVS: LGVS vs. RGVS vs. Sham) × 2 (Velocity: slow vs. fast touch) x× 2 (Order: slow first vs. fast first) mixed ANOVA (with repeated measures on the first two factors) on i) the raw proprioceptive drift values (obtained after each stroking condition), and ii) differential proprioceptive drift scores (calculated by subtracting the drift score obtained during the stroking condition from its immediately preceding visual capture baseline). Paired-samples t-tests (Bonferroni-corrected, α=0.0167) were run to investigate the direction of main effects and interactions. In Experiment 2, we aimed to explore whether seen touch on the Rubber Hand would enhance visual capture effects during LGVS. We hypothesised that such enhancement would be greater for affective rather than neutral touch in the stroking conditions, which was analysed with a mixed 3 (GVS: Sham vs LGVS vs RGVS) × 2 (Velocity: slow vs fast) × 2 (Order: slow first vs fast first) ANOVA, followed by Bonferroni-corrected (α=0.0167) paired-samples t-tests. In both experiments, to investigate subjective feelings of embodiment, we ran all the analyses detailed above on the embodiment questionnaire scores (see SM Sections 2.1.1. and 2.2.1.).

As several of the proprioceptive drift distributions were non-normal, we ran non-parametric analyses to confirm the effects found in the parametric ones (see SM, sections 2.1.4 and 2.2.4).

Data were analysed using the IBM Statistical Package for Social Sciences (SPSS) for Windows, version 23 (Armonk, NY) and plotted using the “*ggplot2*” package for R [40].

## Results

### Experiment 1

#### Visual capture

A 3 (GVS: Sham vs LGVS vs RGVS) × 2 (Order: First vs Second Visual Capture) repeated-measures ANOVA on the proprioceptive drift values of the visual capture conditions revealed a main effect of GVS (F_(2,66)_=6.106, p=.004, ηp^2^=.156) but no main effect of Order (F_(1,33)_=0.114, p=.738, ηp^2^=.003) and no interaction (F_(2,66)_=0.46, p=.955, ηp^2^=.001) (Figure 3A). A subsequent one-way repeated-measures ANOVA on the combined visual capture scores confirmed the main effect of Stimulation (F_(2,66)_=6.11, p=.004, ηp^2^=.156). Planned contrasts (Bonferroni-corrected paired sample t-tests, α=0.025), revealed significantly greater proprioceptive drift following LGVS compared to Sham (LGVS: M=1.75 cm, SD=2.62; Sham: M=0.15 cm, SD=2.16; LGVS vs Sham: t_(33)_=3.410 p=.002 *d*=0.83) and RGVS (RGVS: M=0.50 cm, SD=2.10; LGVS vs RGVS: t_(33)_= 2.459 p=.019 d=0.60) (Figure 3B).

**Figure 3.**
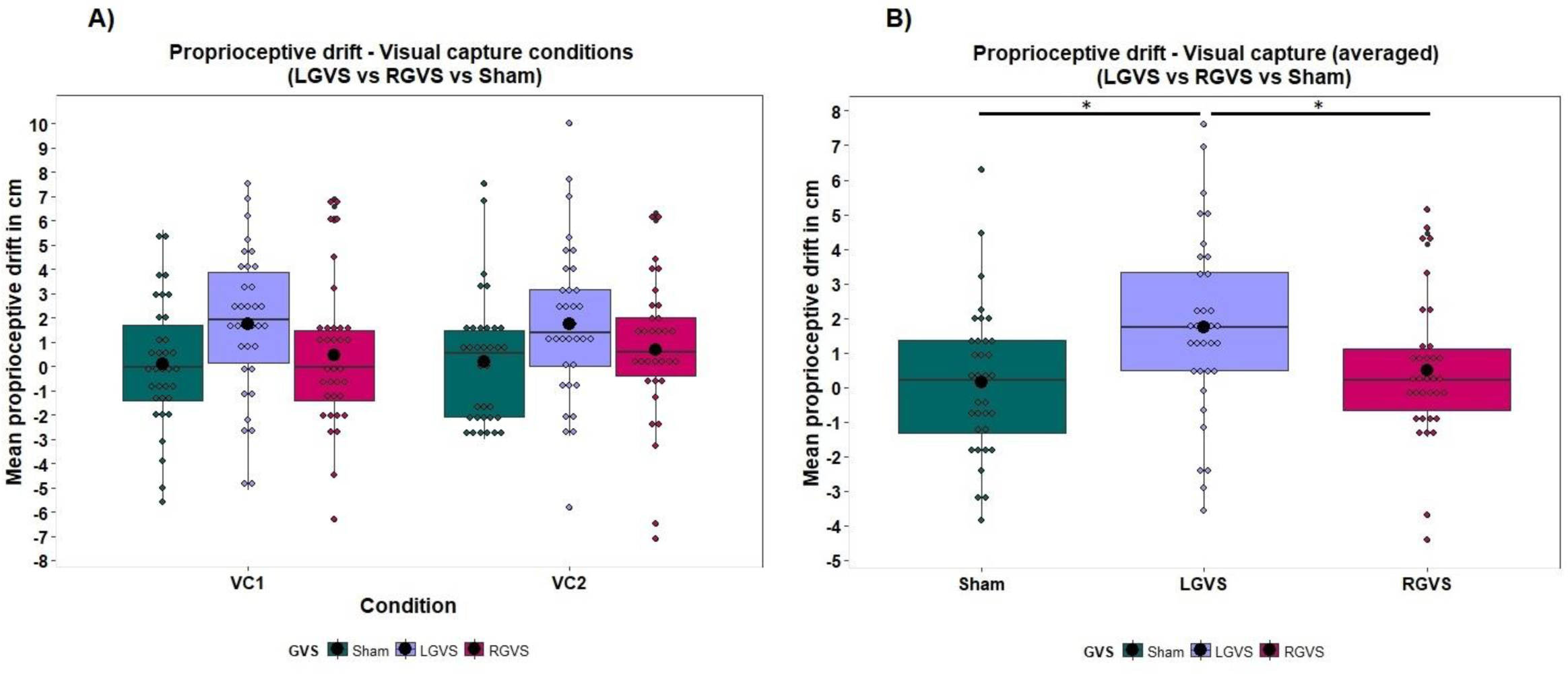
Mean values of the proprioceptive drift measured in cm in Sham, LGVS and RGVS. A) Visual capture conditions 1 and 2 as performed by the participants during the block; B) visual capture baselines averaged. *= p<0.05; Solid line=median; Black dot= mean; Whiskers: upper whisker = min(max(x), Q_3 + 1.5 * IQR); lower whisker = max(min(x), Q_1 – 1.5 * IQR).

#### Stroking conditions

A 3 (GVS: Sham vs LGVS vs RGVS) × 2 (Velocity: Slow vs Fast) × 2 (Order: slow first vs fast first) mixed ANOVA on the proprioceptive drift values obtained from the stroking conditions revealed a significant interaction between Stimulation and Velocity (F_(2,66)_=3.253, p=.045 η_p_^2^=.092) (Figure 4). No main effects or further interactions were significant: Stimulation (F_(2,66)_=880, p=.420, η_p_^2^=.027), Velocity (F _(1,33)_ =1.108, p=.300, η_p_^2^=.033), Order (F _(1,32)_=3.835, p=.059, η_p_^2^=.107), Stimulation*Order (F_(2,64)_=.669, p=.516, η_p_^2^=.020), Velocity*Order (F_(1,32)_=.159, p=.693, η_p_^2^=.005), Stimulation*Velocity*Order (F_(2,64)_=.655, p=.523, η_p_^2^=.020).

**Figure 4.**
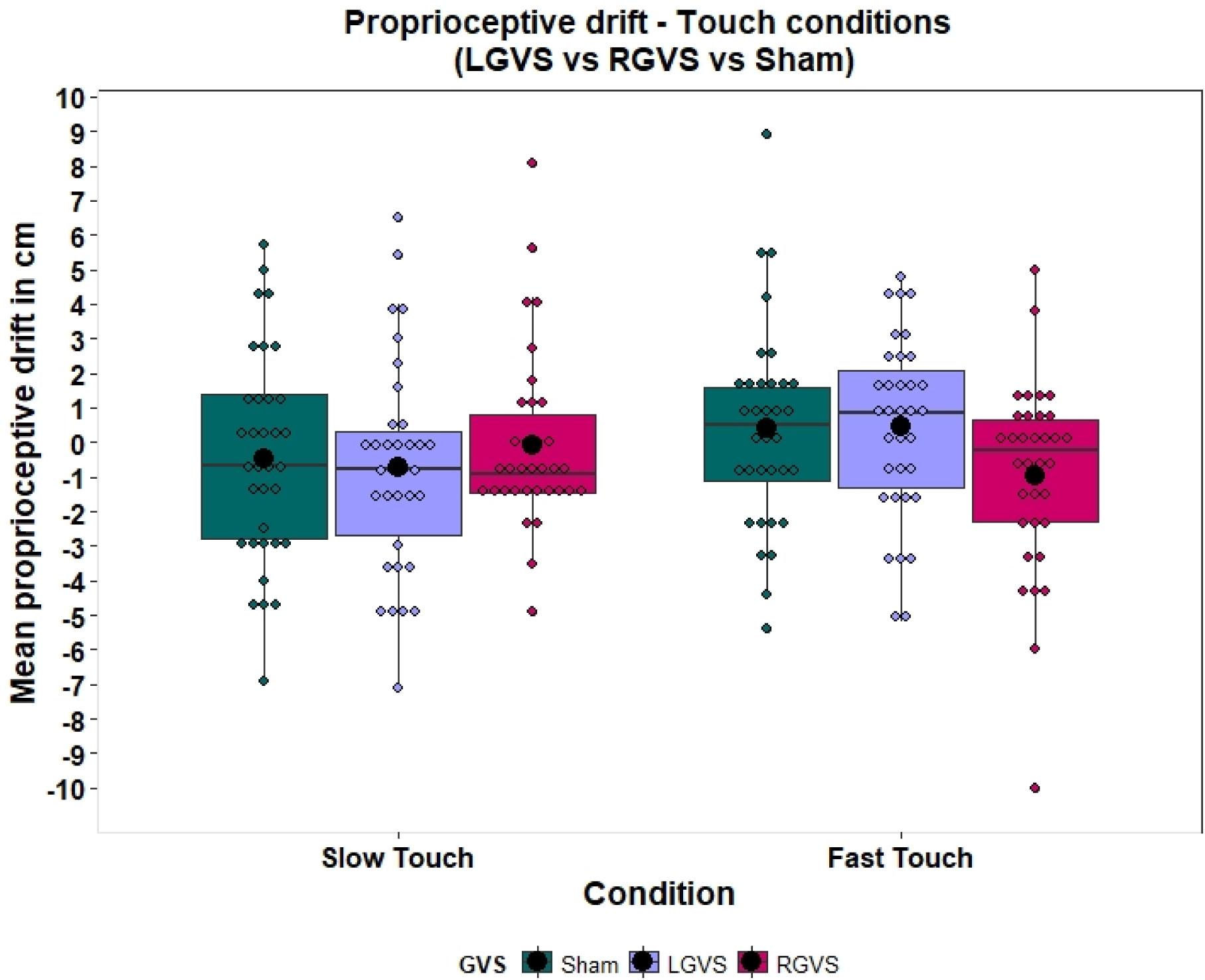
Mean values of the proprioceptive drifts in the different stroking conditions (slow, affective touch and fast, neutral touch) measured in cm and in the three different vestibular configurations (Sham, LGVS and RGVS). Solid line=median; Black dot= mean; Whiskers: upper whisker = min(max(x), Q_3 + 1.5 * IQR); lower whisker = max(min(x), Q_1 – 1.5 * IQR).

To examine the significant interaction, paired-samples t-tests (Bonferroni-corrected, α=0.0167) comparing slow and fast touch for each type of GVS were performed. None of the comparisons were significant (LGVS Slow touch: M=-0.74, SD=3.5; LGVS Fast Touch: M=0.43, SD=2.59; t_(33)_= −2.45, p= 0.020, *d*=0.60; Sham Slow touch: M=-0.46, SD=3.04; Sham Fast Touch: M=0.39, SD=2.92; t_(33)_= −1.323, p= 0.1095, *d*=0.33; RGVS Slow touch: M=-0.08, SD=2.56; RGVS Fast touch: M=-0.96, SD=2.91; t_(33)_=1.287, p= 0.207, *d*=0.32). Non-parametric analysis (section 2.1.4, SM) did not confirm the significant interaction, in line with the non-significant comparisons reported above.

#### Disruption of Visual Capture

A 3-way mixed ANOVA on the differential scores revealed a main effect of Stimulation (F_(2,64)_=4.978, p=.010, η_p_^2^=.135), but no main effect of Velocity (F_(1,32)_=.194, p=.663, η_p_^2^=.006) or Order (F_(1,32)_=.030, p=.864, η_p_^2^=.001, η_p_^2^=.107), and no significant two- or three-way interactions (Figure 5); Vestibular Simulation*Velocity (F_(2,64)_=2.039, p=.139, η_p_^2^=.060), Stimulation*Order (F_(2,64)_=.099, p=.906, ηp^2^=.003), Velocity*Order (F_(1,32)_=.001, p=.975, η_p_^2^=.000), Stimulation*Velocity*Order (F_(2,64)_=.305, p=.738, η_p_^2^=.009). Post-hoc paired samples t-tests comparing each type of GVS regardless of touch velocity (Bonferroni-corrected, α=0.0167) indicated that LGVS led to significantly greater disruption of visual capture (i.e. smaller proprioceptive drifts) in comparison with Sham (LGVS: M=-1.90 cm, SD=2.45; Sham: M=-0.18 cm, SD=2.02; t_(33)_=3.591 p=.001 *d*=0.90) but not RGVS (RGVS: M=-1.02 cm, SD=2.75; t_(33)_=-1.453 p=.156 *d*=0.36), with no difference between RGVS and Sham (t_(33)_=-1.583 p=.123 *d*=0.38).

**Figure 5.**
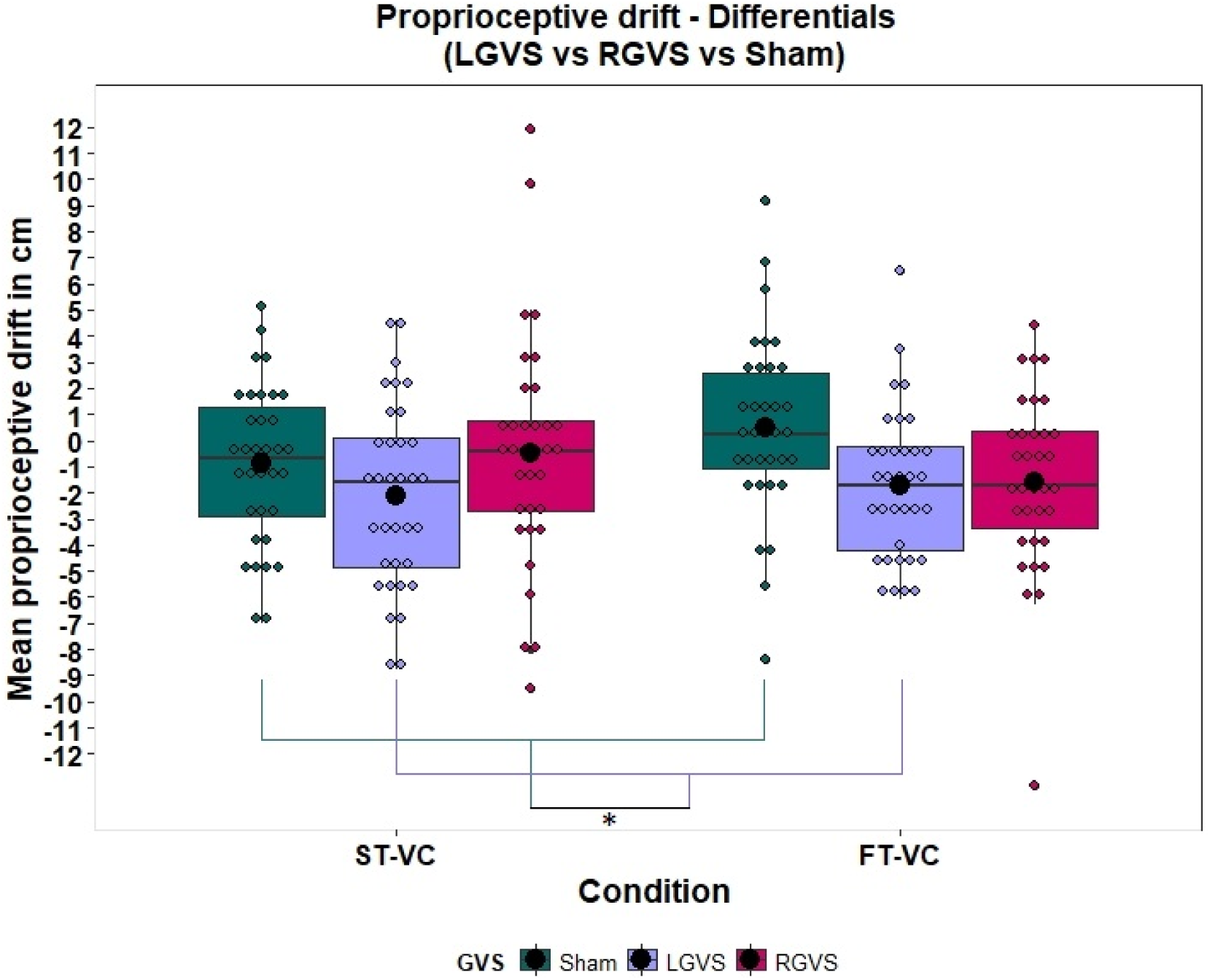
Mean values of the differential scores obtained from the subtraction of the proprioceptive drift values of the visual capture baselines from the stroking conditions’ ones measured in cm and in the three different vestibular configurations (Sham, LGVS and RGVS). ST= slow, affective touch; FT= Fast, neutral touch. *= p<0.05; Solid line=median; Black dot= mean; Whiskers: upper whisker = min(max(x), Q_3 + 1.5 * IQR); lower whisker = max(min(x), Q_1 – 1.5 * IQR).

### Experiment 2

#### Visual capture

A 3 (GVS: Sham vs LGVS vs RGVS) × 2 (Order: First vs Second Visual Capture) repeated-measures ANOVA on the proprioceptive drift values of the two visual capture conditions (Figure 6A), revealed a main effect of Order (F_(1,34)_=5.475, p=.025, ηp^2^=.139), with the first visual capture being higher than the second in all GVS configurations, but no main effect of GVS (F_(2,68)_=1.357, p=.264, ηp^2^=.038) and no interaction (F_(2,68)_=.523, p=.595, ηp^2^=.015). Combining the two visual capture conditions and running a one-way ANOVA comparing GVS blocks revealed the same non-significant results (F_(2,68)_=1.357, p=.264, ηp^2^=.038; Figure 6B). A Friedman’s ANOVA performed on the same data (owing to violation of normality) showed a main effect of stimulation (χ^2^_(2)_=8.647, p=0.013). However, post-hoc Wilcoxon signed-rank tests (α=0.025) were not significant (LGVS: Median=1.25, IQR= 1.85; Sham: Median=0.75, IQR= 1.4; LGVS vs Sham Z=.949 p=0.342; LGVS vs RGVS: RGVS: Median=0.20; IQR= 2.25; LGVS vs RGVS: Z=1.950, p=0.051).

**Figure 6.**
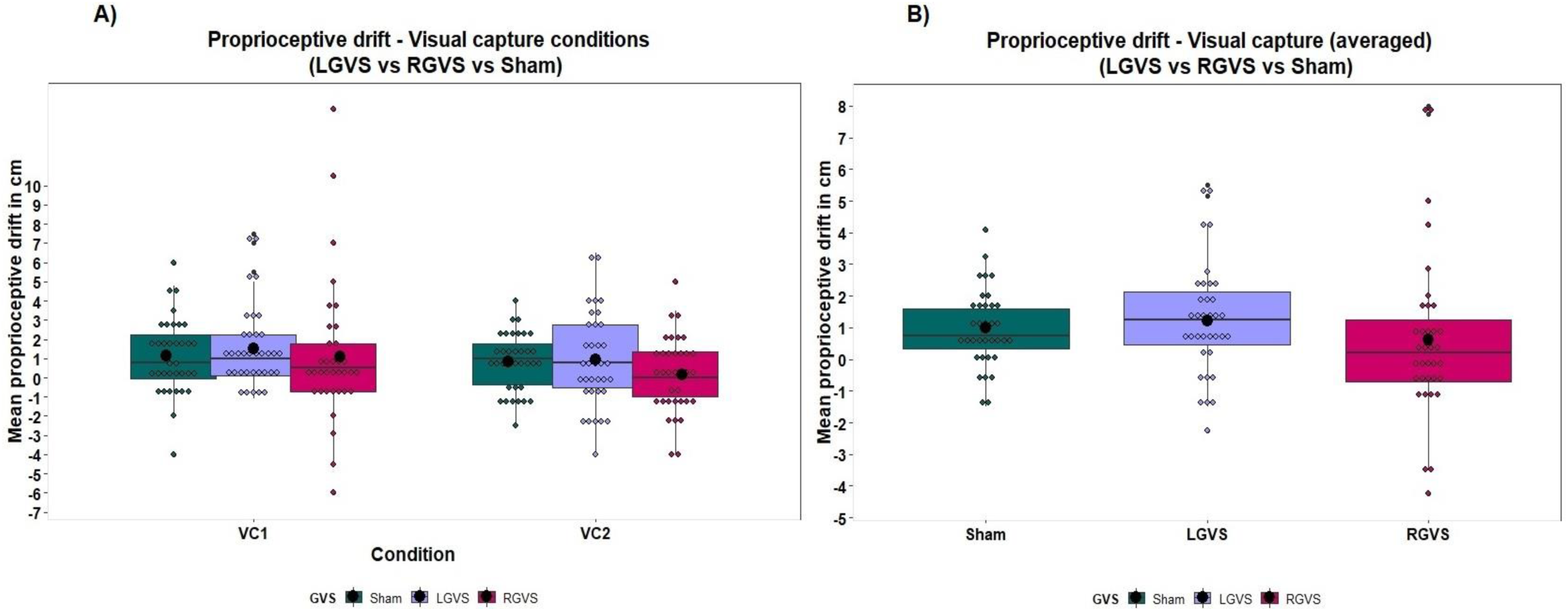
Mean values of the proprioceptive drift measured in cm in Sham, LGVS and RGVS. A) Visual capture conditions 1 and 2 as performed by the participants during the block; B) visual capture baselines averaged. *= p<0.05; Solid line=median; Black dot= mean; Whiskers: upper whisker = min(max(x), Q_3 + 1.5 * IQR); lower whisker = max(min(x), Q_1 – 1.5 * IQR).

#### Visual capture (combined)

To check whether increasing the sample size to include data from both experiments (N=69) would lead to the same results outlined in Experiment 1 and (partly) replicated in Experiment 2, we conducted a 3 (GVS: Sham vs LGVS vs RGVS) x 2 (Order: First vs Second Visual Capture) repeated-measures ANOVA on the proprioceptive drift values of the two visual capture conditions, which revealed a main effect of GVS (F_(2,136)_=5.838, p=.004, ηp^2^=.079) and no Order effects (F_(1,68)_=1.146, p=.288, ηp^2^=.017) or interaction (F_(2,136)_=.134, p=.875, ηp^2^=.002) (Figure 7A). We then averaged the two visual capture conditions and ran a one-way ANOVA on the proprioceptive drifts, confirming such results (GVS: F_(2,136)_=5.954, p=.003, ηp^2^=.081) (Figure 7B). Bonferroni-corrected paired sample t-tests (α=0.025), revealed significantly greater proprioceptive drift during LGVS compared to Sham (LGVS: M=1.48 cm, SD=2.24; Sham: M=0.57 cm, SD=1.79; LGVS vs Sham: t_(68)_=3.221 p=.002 *d*=0.55) and RGVS (RGVS: M=0.56 cm, SD=2.35; LGVS vs RGVS: t_(68)_= 2.818 p=.006 d=0.48). These findings were confirmed by non-parametric tests (SM, section 2.2.4.).

**Figure 7.**
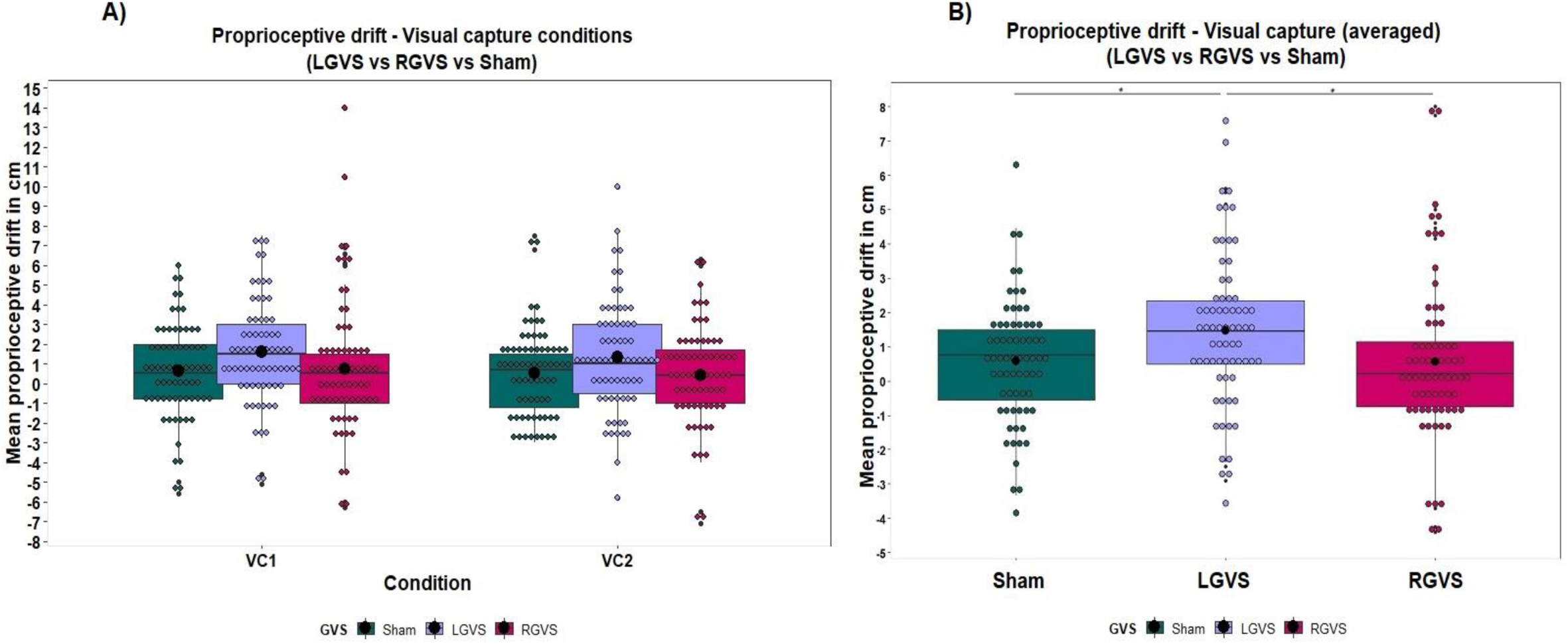
Mean values of the proprioceptive drift measured in cm in Sham, LGVS and RGVS. A) Visual capture conditions 1 and 2 as performed by the participants during the block for both experiment 1 and 2; B) visual capture baselines averaged. *= p<0.05; Solid line=median; Black dot= mean; Whiskers: upper whisker = min(max(x), Q_3 + 1.5 * IQR); lower whisker = max(min(x), Q_1 – 1.5 * IQR).

#### Visual capture of vicarious touch

A 3 (GVS: Sham vs LGVS vs RGVS) × 2 (Velocity: slow vs fast) × 2 (Order: slow first vs fast first) mixed ANOVA on the proprioceptive drift values obtained following the stroking conditions (Figure 8) revealed a main effect of Stimulation (F_(2,66)_=4.608, p=.013, ηp^2^=.123) but no other main effects or interactions; Velocity (F_(1,33)_=.143, p=.708, η_p_^2^=.004), Order (F_(1,33)_=1.986, p=.168, η_p_^2^=.057), Stimulation*Velocity (F_(1,33)_ =.890, p=.416 η _p_^2^=.026), Stimulation*Order (F_(2,66)_ =.245, p=.783, η_p_^2^=.007), Velocity*Order (F_(1,33)_=.001, p=.976, η_p_^2^=.000), Stimulation*Velocity*Order (F_(2,66)_=.878, p=.421, η _p_^2^=.026). Bonferroni-corrected paired samples t-tests (α=0.0167) comparing the effect of GVS irrespective of velocity, indicated that LGVS led to significantly greater visual capture of vicarious touch (i.e. greater proprioceptive drifts) in comparison with Sham (LGVS: M=1.12 cm, SD=1.82; Sham: M=0.33 cm, SD=1.40; t_(34)_=3.187 p=.003 *d*=0.77) but not RGVS (RGVS: M=0.68 cm, SD=1.59; t_(34)_=1.620 p=.114 *d*=0.39) and with no difference between RGVS and Sham (t_(34)_=1.363 p=.182 *d*=0.33).

**Figure 8.**
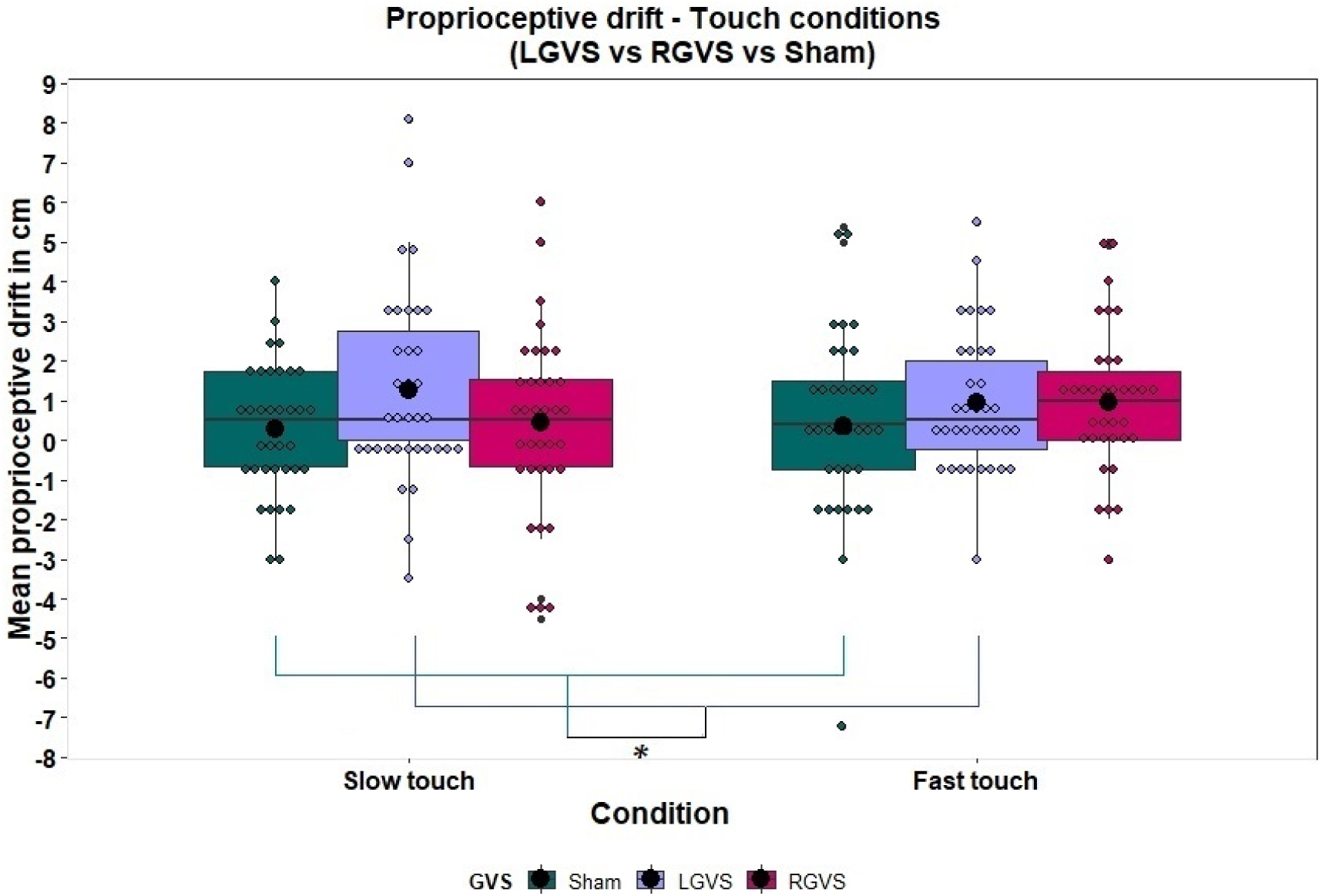
Mean values of the proprioceptive drifts in the different stroking conditions (slow, affective touch and fast, neutral touch) measured in cm and in the three different vestibular configurations (Sham, LGVS and RGVS). Solid line=median; Black dot= mean; Whiskers: upper whisker = min(max(x), Q_3 + 1.5 * IQR); lower whisker = max(min(x), Q_1 – 1.5 * IQR). *p <0.05

## Discussion

We used GVS during an adapted RHI to explore vestibular contributions to multisensory integration aiming to i) replicate our previous findings on visual capture of proprioception (i.e. LGVS leads to greater proprioceptive drifts towards the rubber hand even without touch) and ii) investigate the role of right-hemisphere vestibular stimulation in sensory conflict (i.e. touch felt but not seen and vice-versa). Specifically, we hypothesised that i) LGVS would lead to smaller proprioceptive drifts during tactile stimulation of participants’ skin (i.e. touch felt but not seen) in comparison with RGVS and Sham (“disruption of visual capture”), but in favour of vision when touch is seen but not felt (“visual capture of vicarious touch”) and ii) that both effects would be enhanced by applying affective, slow touch in comparison with neutral, fast touch.

In Experiment 1, we successfully replicated our previous findings: LGVS led to greater visual capture in comparison with Sham and RGVS during mere observation of a rubber hand, i.e. participants showed significantly greater proprioceptive drift towards the rubber hand following right-hemisphere vestibular stimulation. In Experiment 2, we found a similar, yet milder, pattern: LGVS led to greater proprioceptive drifts but not significantly more than RGVS and Sham. This reduction of the effect in Experiment 2 might be due to the different experimental manipulations (felt vs seen touch in the conditions following the visual capture ones) and/or higher individual variability in the sample. However, these findings suggest that stimulation of the right vestibular network may modulate multisensory integration by increasing the weight of vision over proprioception in a visuo-proprioceptive conflict. As we argued elsewhere [23], such visual capture effect may be due to a temporary disruption of participants’ body representation [41,42]. The vestibular system may reduce the relative weight of somatosensory stimuli whilst increasing the relevance of exteroceptive ones in order to allow the resolution of perceptual ambiguity [43]. This would be consistent with the visual capture effects observed in stroke patients with right peri-sylvian lesions [33] and with reports of symptoms remission following right-hemisphere vestibular stimulation in patients with dis-ownership feelings [11,44].

Our second main finding is that vestibular stimulation modulates visuo-tactile conflicts according to whether the touch is felt or seen. In Experiment 1, when touch was applied to participant’s own hand (without concomitant tactile stimulation of the rubber hand), proprioceptive drifts were significantly smaller during LGVS in comparison with Sham stimulation (but not RGVS). In Experiment 2, seen vicarious touch delivered to the rubber hand during LGVS led to increased proprioceptive drifts in comparison with Sham (but nor RGVS). Hence, vestibular signals (not necessarily in a lateralised fashion) may be dynamically contributing to multisensory integration according to the contextual relevance of the different modalities involved. This may explain some of the previous conflicting findings in vestibular stimulation studies (e.g. [21,23] vs [22,45,46]). When a rubber hand is in a plausible position in space, allowing its integration in participants’ body representation (as in our previous and current studies), vestibular signals may contribute to solve perceptual ambiguity by weighting visual signals more than proprioceptive ones. Conversely, when a third sensory modality (i.e. touch) is introduced in an asymmetric fashion, such that incorporation of the rubber hand into participant’s body representation would generate additional conflict (i.e. feeling touch that is not seen leads to increased perceptual ambiguity), vestibular signals do not favour visual cues over proprioceptive ones. However, when touch is seen but not felt (i.e. it is vicariously perceived via vision), vestibular signals seem to favour vision, rather than proprioception, to reduce sensory conflict. Hence, the vestibular system may contribute to the maintenance of a coherent percept of our own body by solving ambiguous perceptual situations: such weighting mechanism could be responsible for the enhancement or reduction of visual cues in visuo-proprioceptive-tactile conflicts according to whether the conflict between the different sensory sources can or cannot be solved via visual dominance over proprioception.

Finally, we did not find differences between affective and neutral touch in disrupting nor enhancing visual capture. This contradicts our hypothesis that the results we observed in our previous study may be due to either the felt or the vicarious properties of affective touch. One possibility is that our previous findings, rather than representing vestibular enhancement of felt or seen components of affective touch, may be explained by the presence of both, delivered in synchrony [47]. Future studies should investigate differential contributions of visuo-tactile versus vicarious and tactile only affective touch to multisensory integration.

To conclude, we provided further evidence that the vestibular system may dynamically contribute to multisensory integration by weighting different sensory modalities according to the context in which they are experienced. In the current study, vestibular stimulation led to an increased dominance of visual information over proprioception during a visuo-proprioceptive conflict as well as during vicarious touch conditions (i.e. when touch was seen on the rubber hand but not felt on participant’s hand), but a decrease of visual capture effects when touch was only felt on participant’s hand but not seen on the rubber hand. These findings suggest that the vestibular network may modulate multisensory experience in a dynamic fashion in an attempt to solve sensory conflicts.

## Supporting information

Experimen 1 - data-set

Experiment 2 - data-set

## Acknowledgements

This work was supported by the European Research Council (ERC) Starting Grant ERC-2012-STG GA313755 (to A.F.) and a University of Hertfordshire studentship (to S.P.).

## Supporting information

Supplementary data can be found online.

## References

[1] Ernst MO, Banks MS. Humans integrate visual and haptic information in a statistically optimal fashion. Nature 2002;415:429–33.

[2] Fetsch CR, Pouget A, DeAngelis GC, Angelaki DE. Neural correlates of reliability-based cue weighting during multisensory integration. Nat Neurosci 2011;15:146–54. doi:10.1038/nn.2983.

[3] Stein BE, Stanford TR, Rowland BA. Development of multisensory integration from the perspective of the individual neuron. Nat Rev Neurosci 2014;15:520–35.

[4] Gallagher S. Philosophical conceptions of the self: implications for cognitive science. Trends Cogn Sci 2000;4:14–21.

[5] Botvinick M, Cohen J. Rubber hands ‘feel’ touch that eyes see. Nature 1998;391:756.

[6] van Beers RJ, Wolpert DM, Haggard P. When feeling is more important than seeing in sensorimotor adaptation. Curr Biol 2002;12:834–7.

[7] Folegatti A, de Vignemont F, Pavani F, Rossetti Y, Farne A. Losing one’s hand: visual-proprioceptive conflict affects touch perception. PLoS One 2009;4:e6920. doi:10.1371/journal.pone.0006920.

[8] Pavani F, Spence C, Driver J. Visual capture of touch: Out-of-the-body experiences with rubber gloves. Psychol Sci 2000;11:353–9.

[9] Rock I, Victor J. Vision and touch: An experimentally created conflict between the two senses. Science 1964;143:594–6.

[10] Been G, Ngo TT, Miller SM, Fitzgerald PB. The use of tDCS and CVS as methods of non-invasive brain stimulation. Brain Res Rev 2007;56:346–61. doi:10.1016/j.brainresrev.2007.08.001.

[11] Bisiach E, Rusconi ML, Vallar G. Remission of somatoparaphrenic delusion through vestibular stimulation. Neuropsychologia 1991;29:1029–31.

[12] Bottini G, Paulesu E, Gandola M, Loffredo S, Scarpa P, Sterzi R, et al. Left caloric vestibular stimulation ameliorates right hemianesthesia. Neurology 2005;65:1278–83.

[13] Brandt T, Dieterich M. The vestibular cortex: its locations, functions, and disorders. Ann N Y Acad Sci 1999;871:293–312.

[14] Cappa S, Sterzi R, Vallar G, Bisiach E. Remission of hemineglect and anosognosia during vestibular stimulation. Neuropsychologia 1987;25:775–82.

[15] Vallar G, Bottini G, Rusconi ML, Sterzi R. Exploring somatosensory hemineglect by vestibular stimulation. Brain 1993;116:71–86.

[16] Lopez C, Schreyer HM, Preuss N, Mast FW. Vestibular stimulation modifies the body schema. Neuropsychologia 2012;50:1830–7. doi:10.1016/j.neuropsychologia.2012.04.008.

[17] Lopez C. The vestibular system: balancing more than just the body. Curr Opin Neurol 2016;29:74–83. doi:10.1097/WCO.0000000000000286.

[18] Ferrè ER, Haggard P. The vestibular body: Vestibular contributions to bodily representations. Cogn Neuropsychol 2016;33:67–81. doi:10.1080/02643294.2016.1168390.

[19] Schmidt L, Artinger F, Stumpf O, Kerkhoff G. Differential effects of galvanic vestibular stimulation on arm position sense in right-vs. left-handers. Neuropsychologia 2013;51:893–9. doi:10.1016/j.neuropsychologia.2013.02.013.

[20] Utz KS, Dimova V, Oppenländer K, Kerkhoff G. Electrified minds: Transcranial direct current stimulation (tDCS) and Galvanic Vestibular Stimulation (GVS) as methods of non-invasive brain stimulation in neuropsychology—A review of current data and future implications. Neuropsychologia 2010;48:2789–810. doi:10.1016/j.neuropsychologia.2010.06.002.

[21] Lopez C, Lenggenhager B, Blanke O. How vestibular stimulation interacts with illusory hand ownership. Conscious Cogn 2010;19:33–47. doi:10.1016/j.concog.2009.12.003.

[22] Ferrè ER, Berlot E, Haggard P. Vestibular contributions to a right-hemisphere network for bodily awareness: combining galvanic vestibular stimulation and the “Rubber Hand Illusion.” Neuropsychologia 2015;69:140–7. doi:10.1016/j.neuropsychologia.2015.01.032.

[23] Ponzo S, Kirsch LP, Fotopoulou A, Jenkinson PM. Balancing body ownership: Visual capture of proprioception and affectivity during vestibular stimulation. Neuropsychologia 2018;117:311–21. doi:10.1016/j.neuropsychologia.2018.06.020.

[24] Craig AD. How do you feel? Interoception: the sense of the physiological condition of the body. Nat Rev Neurosci 2002;3:655–66.

[25] Ceunen E, Vlaeyen JWS, Van Diest I. On the Origin of Interoception. Front Psychol 2016;7. doi:10.3389/fpsyg.2016.00743.

[26] Tsakiris M, Jimenez AT-, Costantini M. Just a heartbeat away from one’s body: interoceptive sensitivity predicts malleability of body-representations. Proc R Soc B Biol Sci 2011;278:2470–6. doi:10.1098/rspb.2010.2547.

[27] Suzuki K, Garfinkel SN, Critchley HD, Seth AK. Multisensory integration across exteroceptive and interoceptive domains modulates self-experience in the rubber-hand illusion. Neuropsychologia 2013;51:2909–17. doi:10.1016/j.neuropsychologia.2013.08.014.

[28] Loken LS, Wessberg J, Morrison I, McGlone F, Olausson H. Coding of pleasant touch by unmyelinated afferents in humans. Nat Neurosci 2009;12:547–8. doi:10.1038/nn.2312.

[29] Crucianelli L, Metcalf NK, Fotopoulou AK, Jenkinson PM. Bodily pleasure matters: velocity of touch modulates body ownership during the rubber hand illusion. Front Psychol 2013;4:703. doi:10.3389/fpsyg.2013.00703.

[30] Crucianelli L, Krahé C, Jenkinson PM, Fotopoulou A (Katerina). Interoceptive ingredients of body ownership: Affective touch and cardiac awareness in the rubber hand illusion. Cortex 2018;104:180–92. doi:10.1016/j.cortex.2017.04.018.

[31] Lloyd DM, Gillis V, Lewis E, Farrell MJ, Morrison I. Pleasant touch moderates the subjective but not objective aspects of body perception. Front Behav Neurosci 2013;7. doi:10.3389/fnbeh.2013.00207.

[32] van Stralen HE, van Zandvoort MJ, Hoppenbrouwers SS, Vissers LM, Kappelle LJ, Dijkerman HC. Affective touch modulates the rubber hand illusion. Cognition 2014;131:147–58. doi:10.1016/j.cognition.2013.11.020.

[33] Martinaud O, Besharati S, Jenkinson PM, Fotopoulou A. Ownership illusions in patients with body delusions: Different neural profiles of visual capture and disownership. Cortex 2017;87:174–85. doi:10.1016/j.cortex.2016.09.025.

[34] Samad M, Chung AJ, Shams L. Perception of body ownership is driven by Bayesian sensory inference. PLoS One 2015;10:e0117178. doi:10.1371/journal.pone.0117178.

[35] Morrison I, Bjornsdotter M, Olausson H. Vicarious Responses to Social Touch in Posterior Insular Cortex Are Tuned to Pleasant Caressing Speeds. J Neurosci 2011;31:9554–62. doi:10.1523/JNEUROSCI.0397-11.2011.

[36] Morrison I, Löken LS, Minde J, Wessberg J, Perini I, Nennesmo I, et al. Reduced C-afferent fibre density affects perceived pleasantness and empathy for touch. Brain 2011;134:1116–26. doi:10.1093/brain/awr011.

[37] Costantini M, Robinson J, Migliorati D, Donno B, Ferri F, Northoff G. Temporal limits on rubber hand illusion reflect individuals’ temporal resolution in multisensory perception. Cognition 2016;157:39–48. doi:10.1016/j.cognition.2016.08.010.

[38] Rohde M, Di Luca M, Ernst MO. The Rubber Hand Illusion: feeling of ownership and proprioceptive drift do not go hand in hand. PLoS One 2011;6:e21659. doi:10.1371/journal.pone.0021659.

[39] Longo MR, Schuur F, Kammers MP, Tsakiris M, Haggard P. What is embodiment? A psychometric approach. Cognition 2008;107:978–98. doi:10.1016/j.cognition.2007.12.004.

[40] ggplot2 - Elegant Graphics for Data Analysis | Hadley Wickham | Springer n.d. https://www.springer.com/gb/book/9780387981413 (accessed February 13, 2019).

[41] Fink GR, Marshall JC, Weiss PH, Stephan T, Grefkes C, Shah NJ, et al. Performing allocentric visuospatial judgments with induced distortion of the egocentric reference frame: an fMRI study with clinical implications. Neuroimage 2003;20:1505–17.

[42] Harris LR, Hoover AEN. Disrupting Vestibular Activity Disrupts Body Ownership. Multisensory Res 2015;28:581–90. doi:10.1163/22134808-00002472.

[43] Zeller D, Litvak V, Friston KJ, Classen J. Sensory Processing and the Rubber Hand Illusion— An Evoked Potentials Study. J Cogn Neurosci 2015;27:573–82. doi:10.1162/jocn_a_00705.

[44] Rode G, Charles N, Perenin M-T, Vighetto A, Trillet M, Aimard G. Partial Remission of Hemiplegia and Somatoparaphrenia Through Vestibular Stimulation in a Case of Unilateral Neglect. Cortex 1992;28:203–8. doi:10.1016/S0010-9452(13)80048-2.

[45] Pavlidou A, Ferrè ER, Lopez C. Vestibular stimulation makes people more egocentric. Cortex 2018;101:302–5. doi:10.1016/j.cortex.2017.12.005.

[46] Pfeiffer C, Serino A, Blanke O. The vestibular system: a spatial reference for bodily self-consciousness. Front Integr Neurosci 2014;8:31. doi:10.3389/fnint.2014.00031.

[47] Filippetti ML, Kirsch LP, Crucianelli L, Fotopoulou A. Affective certainty and congruency of touch modulate the experience of the rubber hand illusion. Sci Rep 2019;9. doi:10.1038/s41598-019-38880-5.

