## Supplementary Materials for "Vestibular modulation of multisensory integration during actual and vicarious tactile stimulation"

1. **Methods**
   1.1. Materials - Embodiment questionnaire

Participants answered the questionnaire [39] using a 7-points Likert scale, ranging from -3 (‘Strongly disagree’) to +3 (‘Strongly agree’) with 0 being ‘Neither agree nor disagree’.

During the block…

Ownership (1-5):

1. …it seemed like I was looking directly at my own hand, rather than at a rubber hand.

2. …it seemed like the rubber hand began to resemble my real hand.

3. …it seemed like the rubber hand belonged to me.

4. …it seemed like the rubber hand was my hand.

5. …it seemed like the rubber hand was part of my body.

Location (6-8):

6. …it seemed like my hand was in the location where the rubber hand was.

7. …it seemed like the rubber hand was in the location where my hand was.

Affect (9):

8. … I found the overall experience enjoyable.

Adapted from the original (“I found that experience interesting”)

1. **Results - Additional analysis**

**2.1. Experiment 1**

2.1.1. Embodiment questionnaire

As described in details in the main Data analysis section, the average values obtained after each condition were analysed as follows. The descriptive statistics and main results are reported in Table 1 below.

**Table 1**. Descriptive statistics of embodiment questionnaire’s average values in the different conditions.

| **Conditions** | **Sham**  Mean *(SD)* | **LGVS**  Mean *(SD)* | **RGVS**  Mean *(SD)* | **ANOVAs** |
| --- | --- | --- | --- | --- |
| Visual Capture (first) | *-0.37* *(1.57)* | -0.56 *(1.73)* | -0.05 *(1.56)* | Main effects*:* Stimulation: F_(2, 66)_=2.196, p=.119, η_p_^2^=.062; Order: F_(1, 33)_=.011, p=.918, η_p_^2^=.000. Interaction: F_(2, 66)_=2.002, p=.143, η_p_^2^=.057 |
| Visual Capture (second) | -0.54 *(1.61)* | -0.30 *(1.71)* | -0.13 *(1.72)* |  |
| Visual Capture (average) | -0.45 *(1.54)* | -0.43 *(1.68)* | -0.10 *(1.54)* | Stimulation: F_(2, 66)_=2.095, p=.131, η_p_^2^=.060 |
| Slow (affective) touch | -0.36 *(1.63)* | -0.53 *(1.75)* | -0.22 (1.65*)* | Main effects: Stimulation: F_(2, 66)_=1.929, p=.153, η_p_^2^=.055; Condition: F_(1, 33)_=.515, p=.478, η_p_^2^=.015. Interaction: F_(2, 66)_=.179, p=.874, η_p_^2^=.005 |
| Fast (neutral) touch | -0.30 *(1.60)* | -0.41 *(1.67)* | -0.19 *(1.51)* |  |
| Differential values (ST-VC) | 0.05 *(0.75)* | -0.08 *(0.82)* | -0.18 *(0.89)* | Main effects: Stimulation: F_(2, 66)_=1.073, p=.348, η_p_^2^=.031; Condition: F_(1, 33)_=2.099, p=.157, η_p_^2^=.060. Interaction: F_(2, 66)_=.120, p=.887, η_p_^2^=.004 |
| Differential values (FT-VC) | 0.20 *(0.79)* | 0.01 *(0.81)* | 0.02 *(0.85)* |  |

Visual capture of proprioception

As for the proprioceptive drift values, to investigate the effects of vestibular stimulation on embodiment, we conducted a 3 (GVS: Sham vs LGVS vs RGVS) x 2 (Order: Visual Capture First vs Visual Capture Second) repeated measures ANOVA on the questionnaire values of the two visual capture conditions. None of the main effects or interactions were significant (Stimulation: F_(2,66)_= 2.196, p=.119, ηp^2^=.062; Order: F_(1,33)_=.011, p=.918, ηp^2^=.000; Stimulation*Order: F_(2,66)_=2.002, p=.143, ηp^2^=.057). To confirm our results, we averaged the two conditions and run an additional one-way repeated measures ANOVA which revealed the same pattern, with no significant main effect of Stimulation: (F_(2,66)_= 2.095, p=.131, ηp^2^=.060).

Stroking conditions

To explore the effects of vestibular stimulation on embodiment following slow, affective touch and fast, neutral touch, we conducted a 3 (GVS: Sham vs LGVS vs RGVS) x 2 (Velocity: Slow touch vs Fast touch) repeated measures ANOVA on the questionnaire values of the two stroking conditions. None of the main effects or interactions were significant (Stimulation: F_(2,66)_=1.929, p=.153, ηp^2^=.055; Velocity: F_(1,33)_=.515, p=.478, ηp^2^=.015; Stimulation*Velocity: F_(2,66)_=.179, p=.874, ηp^2^=.005).

Differentials

To investigate the effects of vestibular stimulation and touch in disrupting visual capture, we calculated a differential value by subtracting the scores obtained following each visual capture condition from the ones obtained after the subsequent stroking condition (see Data analysis and Results sections above for further details). We conducted a 3 (GVS: LGVS vs RGVS vs Sham) x 2 (Velocity: Slow touch vs Fast touch) repeated measures ANOVA, revealing no main effect or interaction (Stimulation: F_(2,66)_=1.073, p=.348, ηp^2^=.031; Velocity: F_(1,33)_=2.099, p=.157, ηp^2^=.060; Stimulation*Velocity: F_(2,66)_=.120, p=.887, ηp^2^=.004).

2.1.2. Enjoyability

One of the statements of the embodiment questionnaire (i.e. 8. “… I found the overall experience enjoyable.”) capturing affective aspects of the experience was analysed separately. The same analyses described above for the embodiment questionnaire were implemented and none of them showed significant results.

2.1.3 Cumulative effects of GVS

In this analysis, we aimed to explore whether vestibular stimulation had any cumulative, carry over effects on proprioception, over and above any effects of our task manipulations. We analysed the proprioceptive judgements taken before each of the stroking conditions and investigated their trend in each of the three GVS blocks (Sham, LGVS, RGVS) and compared across them. We conducted a 3 (stimulation: LGVS, RGVS and Sham) x 4 (time: 1st Proprioceptive judgement, 2nd proprioceptive Judgement, 3rd Proprioceptive Judgement, 4th Proprioceptive judgement) repeated measures ANOVA, that revealed a main effect of Time (F(_3,99_)=10.319, p<0.001, ηp^2^=.238), indicating a progressive increase in proprioceptive displacement during each block, but no main effect of Stimulation (F(_2,66_)=1.403, p=.253, ηp^2^=.041) nor significant interaction between Stimulation and Time (F(_6,198_)=1.391, p=.220, ηp^2^=.040). An analysis of the polynomial contrasts revealed a significant linear (F(_1,33_)=15.801, p<0.001, ηp^2^=.324) as well as cubic (F(_1,33_)=9.254, p<0.01, ηp^2^=.219) trend over time, suggesting that regardless of the type of stimulation involved, participants’ proprioceptive judgements increased following visual capture conditions and decreased following touch ones (Supplementary Figure 1). This analysis suggests that our findings concerning the proprioceptive drifts (i.e. the differentials between post and pre proprioceptive judgements) were not biased by a cumulative carry-over effect of vestibular stimulation on baseline proprioceptive judgements.

Pre-condition proprioceptive judgements

Proprioceptive judgements

|  | Sham | LGVS | RGVS |
| --- | --- | --- | --- |
| VC1 | 3.09706 | 1.22353 | 1.69706 |
| Touch condition 1 | 3.56471 | 3.37647 | 3.59412 |
| VC 2 | 3.12941 | 2.86177 | 2.73529 |
| Touch condition 2 | 3.93235 | 3.97941 | 4.16765 |

**Supplementary figure 1.** Pre-condition proprioceptive judgements in Sham, LGVS and RGVS across each GVS block (i.e. values taken from the four conditions in the order they were performed). Error bars: SEM.

2.1.4. Non parametric analysis

Data analysis

Main effects of Stimulation were analysed by calculating the average of the proprioceptive drifts or questionnaire values obtained in each of the vestibular stimulations regardless of the order of the condition and comparing these values using Friedman’s ANOVA. The same procedure was followed in order to check for main effects of Order of the conditions, i.e. we averaged the values across the three GVS configurations and compared the results obtained via a Wilcoxon signed rank test.

2-way interactions (Stimulation*Order) were obtained by subtracting one level of the factor of interest from the other one (e.g. for the interaction between Stimulation and Stroking condition, the subtraction has been performed between LGVS slow, affective touch and fast, neutral touch and the resulting distribution has then been compared with the equivalent subtraction in Sham and RGVS). The results were then analysed via a Wilcoxon signed rank test.

As mentioned above, three of the proprioceptive drifts distributions were non normal, according to Shapiro-Wilk tests (Visual capture one - Sham: S-W: 0.896, p=0.004; Slow Touch RGVS: S-W: .879, p=0.001; Fast Touch RGVS: S-W: .937 p=0.049). Hence, we ran non-parametric analyses on the interested proprioceptive drift distributions in order to confirm the effects found in the parametric analysis reported in the Results section above.

Proprioceptive drift

Visual capture of proprioception

Given that one of the two visual capture distribution in Sham was non normally distributed, we run a non-parametric equivalent of the ANOVA reported in the main results section in order to check for main effects of order of the conditions. This analysis confirmed our findings, i.e. there is no significant difference between the two conditions (Z= 0.222, p=0.824).

Stroking conditions

In order to examine the effects of vestibular stimulation on proprioceptive drifts following stroking, we ran a Friedman’s ANOVA on the averaged values of the two stroking conditions, which did not show any difference between the three GVS configurations (Z=3.173, p=0.205). We then averaged the proprioceptive drift values across the different GVS configurations and used a Wilcoxon signed-rank test in order to check for differences due to the stroking conditions, which did not yield significant results (Z=1.402, p=0.161). The interaction between the two factors was tested by subtracting the values of fast touch from the slow touch ones within each GVS configuration and then comparing them via a Friedman’s ANOVA, which did not reveal a difference between the conditions (Z=1.111, p=0.574).

2.1.5. Manipulation checks

Pleasantness task: design and measures

In a separate task at the end of the main experiment we aimed to check whether or not slow touch was perceived as more pleasant than fast touch. Participants were asked to rate the pleasantness of stroking received at 3 and 18 cm/s, in a 2 (velocities: slow vs. fast) x 2 (site: forearm vs. palm of the hand) within-subjects design. Touch was delivered either on the forearm (i.e. a CT site) or on the palm of the hand (i.e. non CT site) [31]. There were four different conditions in total: slow touch on the forearm, slow touch on the palm, fast touch on the forearm and fast touch on the palm.

Two adjacent stroking areas of 6x4 cm each were drawn with a washable marker on the participant’s left forearm and palm of the left hand. Participants were instructed to verbally report, with their eyes closed, the pleasantness of the touches received either on the palm or on the forearm, on a 0 (‘Not at all pleasant’) to 100 (‘Extremely pleasant’) scale. In total, there were 12 trials: 3 trials per each combination of velocities and sites. All the touches were delivered using the same brush described above. During the slow touches, each area was stroked for 2 seconds (at a rate of 3 cm/s) with a 1 second interval in between the 2 strokes, whereas during the fast brushes each area was stroked for 0.3 seconds. The order of the trials was randomised and there was a short break in between each trial, during which participants reported their pleasantness ratings.

We ran a 2 (Velocity: Slow touch vs. Fast touch) x 2 (Site: Forearm vs. Palm of the hand) repeated measures ANOVA on the pleasantness ratings, which revealed that Slow touch was perceived as more pleasant than Fast touch (Velocity: F_(1, 33)_=38.205, p<0.001, η^2^=.537; Slow touch: mean: 78.28 SD: 14.69; Fast touch: mean: 62.75; SD: 18.85) with no difference between the two sites (Site: F_(1, 33)_=.252, p=.619, η^2^=.008) and no interaction between Velocity and Site (F_(1, 33)_=1.268, p=.268, η^2^=.037). Given that Slow touch ratings on the palm were not normally distributed (S-W: .924, p=.021), we calculated main effects and differential scores (using the methods described above in the “non-parametric analysis – data analysis” section) and subsequently ran non-parametric analysis using Wilcoxon-signed rank tests. The results confirmed the findings of the parametric analysis (Velocity: Z=- 4.779, p=0.000; Slow touch: median: 81.67 IQR: 20; Fast touch: median: 64.58; IQR: 22.46; Site: Z=.206, p=.837; Velocity*Site: Z=.535, p=.593)

Vestibular-induced sensations

At the end of the main experiment, participants were asked to report any physical sensation associated with the stimulation in order to identify any noticeable vestibular-induced sensations between conditions. The majority of the sample (32/35) reported a tingling or itching sensation under the patches that none of the participants described as painful. About 2/3 of the sample (23/36 participants) reported vestibular-related sensations (dizziness, vertigo or loss of balance). This suggests that a 1mA stimulation has been able to trigger distinctive vestibular sensations in two thirds of the sample, in line with previous findings from our group [23].

In order to control for the role of Sham stimulation as a placebo-like intervention, we asked the participants to guess in which of the 3 GVS configurations they thought to have received a vestibular system’s stimulation, specifying that the answer might have been all of them, 2 or 1. The majority of them guessed correctly about LGVS and RGVS (31/36 and 30/36, respectively), whereas two thirds of the sample reported Sham as a vestibular stimulation (24/36) when directly invited to compare it with the other 2 GVS configurations.

**2.2. Experiment 2**

2.2.1. Embodiment questionnaire

Data analysis

As described in details in the main Data analysis section above, the average values obtained after each condition were analysed as follows (see Table 2 for descriptive statistics and a summary of main results).

**Table 2**. Descriptive statistics of embodiment questionnaire’s average values in the different conditions.

| **Conditions** | **Sham**  Mean *(SD)* | **LGVS**  Mean *(SD)* | **RGVS**  Mean *(SD)* | **ANOVAs** |
| --- | --- | --- | --- | --- |
| Visual Capture (first) | -.15 *(1.57)* | -*.08* *(1.77)* | -.03 *(1.72)* | Main effects*:* Stimulation: F_(2,68 )_=.513, p=.601, η_p_^2^=.015; Order: F_(1, 34)_=.208, p=.652, η_p_^2^=.006. Interaction: F_(2, 68)_=.171, p=.843, η_p_^2^=.005. |
| Visual Capture (second) | -.11 *(1.70)* | -.09 *(1.68)* | -.07 *(1.65)* |  |
| Visual Capture (average) | -.13 *(1.55)* | -.09 *(1.69)* | -.02 *(1.65)* | Stimulation: F_(2, 68)_=.524, p=.595, η_p_^2^=015. |
| Slow (affective) touch | -.06 *(1.60)* | .12 *(1.62)* | -.04 (1.75*)* | Main effects: Stimulation: F_(2, 68)_=1.244, p=.295, η_p_^2^=.035; Condition: F_(1, 34)_=.399, p=.532, η_p_^2^=.012. Interaction: F_(2, 68)_=.394, p=.676, η_p_^2^=.011 |
| Fast (neutral) touch | -.12 *(1.63)* | -.01 *(1.75)* | -.03 *(1.78)* |  |

Visual capture of proprioception

To explore the effects of vestibular stimulation on embodiment, we conducted a 3 (GVS: Sham vs LGVS vs RGVS) x 2 (Order: Visual Capture First vs Visual Capture Second) repeated measures ANOVA on the questionnaire values of the two visual capture conditions. None of the main effects or interactions were significant (Stimulation: F_(2,68)_=.513, p=.601, ηp^2^=.015; Order: F_(1,34)_=.208, p=.652, ηp^2^=.006; Stimulation*Order: F_(2,68)_=.171, p=.843, ηp^2^=.005). To confirm our results, we averaged the two conditions and run an additional one-way repeated measures ANOVA which showed no significant main effect of Stimulation: (F_(2,68)_=.524, p=.595, ηp^2^=.015).

Stroking conditions

To explore the effects of vestibular stimulation on embodiment following slow, affective and fast, neutral vicarious touch on the rubber hand, we conducted a 3 (GVS: Sham vs LGVS vs RGVS) x 2 (Velocity: Slow touch vs Fast touch) repeated measures ANOVA on the questionnaire values of the two stroking conditions. None of the main effects or interactions were significant (Stimulation: F_(2,68)_=1.244, p=.295, ηp^2^=.035; Velocity: F_(1,34)_=.399, p=.532, ηp^2^=.012; Stimulation*Velocity: F_(2,68)_=.394, p=.676, ηp^2^=.011).

2.2.2. Enjoyability

One of the statements of the embodiment questionnaire (i.e. 8. “… I found the overall experience enjoyable.”) capturing affective aspects of the experience was analysed separately. As for Experiment 1, the same analyses described above for the embodiment questionnaire were implemented, with none of them yielding significant results.

2.2.3. Cumulative effects of GVS

As for Experiment 1, we aimed to confirm that vestibular stimulation did not have any cumulative, carry over effects on proprioception, on the top of the effects of our manipulations. We analysed the proprioceptive judgements taken before each of the stroking conditions and analysed their trend in each of the three GVS blocks (Sham, LGVS, RGVS) and compared across them. We ran a 3 (Stimulation: LGVS, RGVS and Sham) x 4 (time: 1st Proprioceptive judgement, 2nd proprioceptive Judgement, 3rd Proprioceptive Judgement, 4th Proprioceptive judgement) repeated measures ANOVA, which did not reveal any significant main effect or interaction (Stimulation: (F(_2,68_)=.378, p=.687, ηp^2^=.011; Time: F(_3,102_)=.056, p=983, ηp^2^=.002; Stimulation*Time: F(_6,204_)=.581, p=.745, ηp^2^=.017). This analysis suggests that our findings were not biased by a cumulative carry-over effect of vestibular stimulation on baseline proprioceptive judgements nor on a cumulative effect of the illusion (conversely to what we found in Experiment 1).

2.2.4. Non parametric analysis

Data analysis

Since seven of the proprioceptive drifts distributions were non normal, according to Shapiro-Wilk tests (Slow Touch LGVS: S-W: .934 , p=0.037; Visual capture two - LGVS: S-W: 0.877, p=0.001; Visual capture (first) – LGVS: S-W: .870, p=0.001; Visual capture one - RGVS: S-W: .822, p<0.001; Fast Touch RGVS: S-W: .879, p=0.001; Visual capture (average) - RGVS: S-W: 885, p=0.002; Visual capture (first) – RGVS: S-W: .847, p<0.001.), we ran non-parametric analyses on the interested proprioceptive drift distributions in order to confirm the effects found in the parametric analysis reported in the Results section. The procedure implemented to obtain the data and the analyses conducted are the same as those explained above for experiment 1.

Proprioceptive drift

Visual capture of proprioception

The main effect of order revealed by the repeated measures ANOVA reported in the main results section was confirmed by a Wilcoxon signed-rank test (Z= 2.039, p=0.041). The findings on the main effect of stimulation are reported in the main results section.

Visual capture (combined samples)

To confirm the findings of our exploratory analysis on the visual capture data from both Experiment 1 and 2, we ran a Friedman’s ANOVA on the average of the two visual capture conditions, revealing a main effect of stimulation (χ^2^_(2)_ =19.737, p<0.0001), further investigate via post—hoc Wilcoxon signed-rank test (Bonferroni corrected, α= 0.025). Such comparisons showed that LGVS led to greater proprioceptive drifts in comparison with both Sham (Z= 3.121; p=0.002) and RGVS (Z= 2.822; p=0.005).

Stroking conditions

In order to examine the effects of vestibular stimulation on proprioceptive drifts following seen touch, we conducted a Friedman’s ANOVA on the averaged values of the two stroking conditions, which confirmed the parametric analysis reported above (χ^2^_(2)_ = 7.667, p=0.022). To explore this main effect of stimulation, we ran post-hoc Wilcoxon signed-rank test (Bonferroni corrected, α=0.0167), confirmed the results reported above, i.e. that LGVS increased proprioceptive drifts in comparison with Sham (Z=2.786, p= 0.005) but not RGVS (Z=1.223, p=0.221), with no difference between Sham and RGVS (Z=1.172 , p=0.241) and regardless of the type of touch.

2.2.5. Manipulation checks

Pleasantness task: design and measures

Both methods and parametric data analysis were the same as the ones described for experiment 1 for the pleasantness task. In addition to the above, we added an extra block of 6 trials (3 of which included administration of 2 slow, affective consecutive strokes and 3 entailing 2 fast, neutral strokes) in which we administered touch only to the rubber hand while it was in full view and we asked participants to rate how pleasant they thought the touch would be (on the same 0-100 rating scale described above). This block of vicarious pleasantness ratings was counterbalanced across participants (i.e. half the participants did the vicarious rubber hand block first whilst the other half started with the actual touch block), in order to avoid the influence of perceiving touch on participants’ own skin on vicarious ratings.

On the actual touch block, a 2 (Velocity: Slow touch vs. Fast touch) x 2 (Site: Forearm vs. Palm of the hand) repeated measures ANOVA on the pleasantness ratings was conducted, which confirmed that Slow touch was perceived as more pleasant than Fast touch (Velocity: F_(1, 34)_=24.518, p<0.001, η^2^=.491; Slow touch: M=70.67, SD=14.56; Fast touch: M=61.5, SD=15.26) with no difference between the two sites (Site: F_(1, 34)_=2.371, p=.133, η^2^=.065) and no interaction between Velocity and Site (F_(1, 34)_=.036, p=.850, η^2^=.001). A paired-samples t-test was run on the vicarious rubber hand touch block, revealing that participants rated slow touch as more pleasant than fast touch (Slow touch: M=69.4, SD=18.84; Fast touch: M=58.78, SD=15.99; t_(34)_=5.154 p<0.001 *d*=1.25) even when touch was only seen but not felt.

Finally, we run a 2 (Felt touch: slow vs fast) x 2 (Vicarious touch: slow vs fast) repeated-measures ANOVA to check differences between slow and fast touch in the separate felt and vicarious block. In this analysis we averaged across sites for the felt block as there were no differences between forearm and palm. Results revealed a main effect of Velocity (F_(1, 34)_=28.407, p<0.001, η^2^=.455; Slow touch: M=70.03, SD=16.73; Fast touch: M=60.15; SD=15.58) but no differences between the blocks (Type of block: F_(1, 34)_=.445, p=.509, η^2^=.013) and no interaction between velocity and type of block (F_(1, 34)_=1.370, p=.250, η^2^=.039). The addition of the vicarious Rubber Hand block confirmed that, as suggested by previous research [35,36], participants perceived touch administered on the rubber hand significantly more pleasant when it was slow compared with fast. Additionally, the comparison between the pleasantness reported for the felt and the seen touch did not differ and only highlighted that participants rated affective rather than neutral touch as more pleasant regardless of whether it was felt or seen (Supplementary Figure 2).

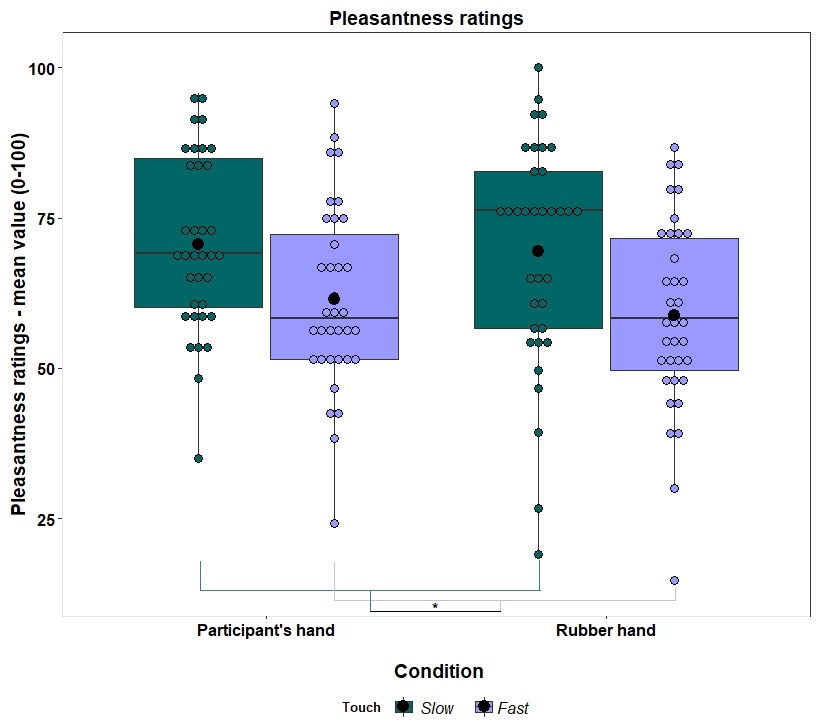

**Supplementary Figure 2.** Mean values of the pleasantness ratings in the felt vs seen touch conditions according to the touch velocity (slow, affective vs fast, neutral). Ratings were on a scale from 0 “Not at all pleasant” to 100 “Extremely pleasant”. Solid line=median; Black dot= mean; Whiskers: upper whisker = min(max(x), Q_3 + 1.5 * IQR); lower whisker = max(min(x), Q_1 – 1.5 * IQR). *p <0.001.

Vestibular-induced sensations

At the end of the main experiment, participants were asked to report any physical sensation associated with the stimulation in order to identify any noticeable vestibular-induced sensations between conditions. The majority of the sample (25/35) reported a tingling or itching sensation under the patches that none of the participants described as painful. About 1/3 of the sample (14/36 participants) reported vestibular-related sensations (dizziness, vertigo or loss of balance), suggesting that a 1mA stimulation has been able to trigger distinctive vestibular sensations in 1/3 of the sample, partially in line with previous findings.

To control for the role of Sham as a placebo-like intervention, we asked participants to guess in which of the 3 GVS configurations they thought to have received a vestibular stimulation, specifying that the answer might have been all of them, 2 or 1. The majority guessed correctly about LGVS and RGVS (28/35 and 19/35, respectively), whereas approximately 1/3 of the sample reported Sham as a vestibular stimulation (13/35) when directly invited to compare it with the other 2 GVS configurations.
